# A geometric deep learning framework for drug repositioning over heterogeneous information networks

**DOI:** 10.1101/2022.07.16.500310

**Authors:** Bo-Wei Zhao, Xiaorui Su, Peng-Wei Hu, Yu-Peng Ma, Xi Zhou, Lun Hu

## Abstract

The effectiveness of computational drug repositioning techniques has been further improved due to the development of artificial intelligence technology. However, most of the existing approaches fall short of taking into account the non-Euclidean nature of biomedical data. To overcome this problem, we propose a geometric deep learning (GDL) framework, namely DDAGDL, to predict drug-disease associations (DDAs) on heterogeneous information networks (HINs). DDAGDL can take advantage of complicated biological information to learn the feature representations of drugs and diseases by ingeniously projecting drugs and diseases including geometric prior knowledge of network structure in a non-Euclidean domain onto a latent feature space. Experimental results show that DDAGDL is able to identify high-quality candidates for Alzheimer’s disease (AD) and Breast neoplasms (BN) that have already been reported by previously published studies, and some of them are not even identified by comparing models.

## Introduction

Traditional drug research and development is a long process mean that requires higher costs and its benefits are estimated that less than a dollar after return on average. Unfortunately, very few drugs end up on the market not even including when the drug was approved [1]. However, various rare diseases are still on the rise, to threaten human health. Take AD as an example, more than 40 million people have AD worldwide, and it will increase even more, but no specific medications have been licensed for use in individuals with mild cognitive impairment [2].

Drug repositioning (also called drug repurposing) is a promising strategy to discover new indicators for approved or experimental drugs, which offers vast advantages to accelerating the development of new drug [3]. In recent years, many computational-based methods for discovering new indicators of approved drugs have been developed to improve the efficiency of drug discovery and development [4-8]. Generally, these approaches for in silico drug repositioning can be classified into three categories, including network-based [9, 10], matrix factorization-based [11, 12], and deep learning-based approaches [13, 14].

Network-based approaches predict unknown DDAs by learning characteristics of drugs and diseases from integrated multiple drug-related networks. For instance, deepDR [10] first integrates a heterogeneous network that includes drug-disease, drug-side-effect, drug-target and seven drug-drug networks, and then obtains the features of drugs and diseases by a multi-modal deep autoencoder to fuse each network information, and finally uses a variational autoencoder to infer indicators for approved drugs. Matrix factorization-based approaches can decompose the high-dimensional association matrix into the product of two low-dimensional matrices to recommend candidates for diseases. DTINet [12] obtains low-dimensional vector representations by a compact feature learning algorithm from a heterogeneous network with a variety of drug-related networks and then discovers new interactions for drugs and targets. However, these approaches generally fail to take into account the non-Euclidean nature of biomedical data to capture more impactful features for DDA prediction, and then the features propagated through biological networks are more susceptible to biological association network data and noise.

Recently, deep learning approaches have been particularly successful when dealing with biological data with underlying Euclidean structure [6, 8, 15-19]. As more and more biological data are discovered, these biological data not only include invariant biological attributes, such as the amino acid sequence of proteins, the base sequence of RNA molecules and the molecular structure of drugs, but also their network structure information should be considered, i.e. non-Euclidean nature. However, this information cannot be computed by previous deep learning. Therefore, geometric-based deep learning techniques, which can capture the features of biological data with non-Euclidean nature by projecting these data into a latent feature space, are starting to receive more attention. DRHGCN [14] adopts multiple graph convolutional layers to capture the embedding representations of drugs and diseases from three networks including the drug-disease, drug-drug similarity and disease-disease similarity networks. Although effective, it is limited by the over-smoothing of the graph convolutional network, and it is difficult to fully capture the feature representation of drugs and diseases for a more accurate predicting DDAs.

In this paper, a novel drug repositioning framework, called DDAGDL, is developed by using geometric deep learning in a HIN. DDAGDL can not only cope with non-Euclidean data and high-dimensional biological association data from biological heterogeneous networks, but also select the optimal feature space to improve the expression ability of feature representations of drugs and diseases. DDAGDL projects the biomedical data with non-Euclidean nature into the latent feature space to capture the feature representation of each biomedical molecule (i.e., drugs, proteins and disease) across multiple biological networks. Based on geometric deep learning, DDAGDL then judges an optimal projection for drugs and diseases by multiple neural network propagation. Experiment results on three benchmark datasets demonstrate the superior performance of DDAGDL when comparing it with several state-of-the-art drug repositioning models. Furthermore, we have also conducted the case studies to show the usefulness of DDAGRL in predicting novel DDAs by validating the top-ranked drug candidates predicted by DDAGRL for AD and BN. Our findings indicate that most of drug candidates are with high quality, as they have already reported by previously published studies, and some of them are not even found in the prediction results of the other comparing models. In this regard, leveraging geometric deep learning provides us an alternative view to address the problem of drug repurposing by properly handling the non-Euclidean nature of biomedical data, which has been ignored by most of existing prediction models. In conclusion, we believe that our work opens a new avenue in drug repositioning with new insights gained from geometric deep learning.

## Results

### Overview of DDAGDL

DDAGDL is composed of three steps. Frist, DDAGDL calculates the biological attribute of all biomedical data in the HIN. Second, three biomedical molecules are projected into the latent feature space according to the biological attribute and the geometric prior knowledge of network structure in a non-Euclidean domain to further obtain more influential feature representations for drugs and diseases, in which DDAGDL judges the best projection space for drugs and diseases by multiple neural network propagation. After that, DDAGDL infers new interactions between drugs and diseases by the scores predicted of the XGBoost classifier.

### Comparison with state-of-the-art drug repositioning models

To accurately evaluate the performance of DDAGDL, we first use a ten-fold cross-validation (CV) scheme. In particular, a benchmark dataset is divided into 10 subsets, each subset is alternatively taken as a testing set while the remaining subsets as the training set, in which randomly sampled non-interacting pairs that the number of matches equal to the known drug-disease pairs is held out as negative samples. In addition, the experimental results are shown in the Supplementary material. More importantly, we have compared DDAGDL with three state-of-the-art models for drug repositioning, including deepDR [10], DTINet [12], and DRHGCN [14]. A variant of DDAGDL, i.e., DDAGDL-A, is implemented, which only considers the biological attribute of drugs and diseases, to study the influence of the geometric deep learning strategy for identifying the relationships between biomedical entities.

Regarding the setting of parameters involved when training these drug repositioning models on three benchmark datasets, we adopt the default parameter settings for the competing models, i.e., deepDR, DTINet, and DRHGCN, as recommended in their public codes for a fair comparison.

The experimental results of 10-fold CV on B-dataset, C-dataset and F-dataset are presented in Tables 1, 2 and 3, and Figure 2. We note that DDAGDL surpasses the all-comparison algorithms across three benchmark datasets in terms of ACC, MCC, F1-score, and AUC. In this regard, we guess this is a strong indicator for applying to the discovery of new indications, due to DDAGDL is preferred over state-of-the-art models. The comparison results show that DDAGDL performs a superior performance in terms of the average AUC across three benchmark datasets, as it has better by 6.10%, 2.54%, 7.61% and 5.00% than deepDR, DRHGCN, DDAGDL-A and DTINet, respectively.

**Table 1.**
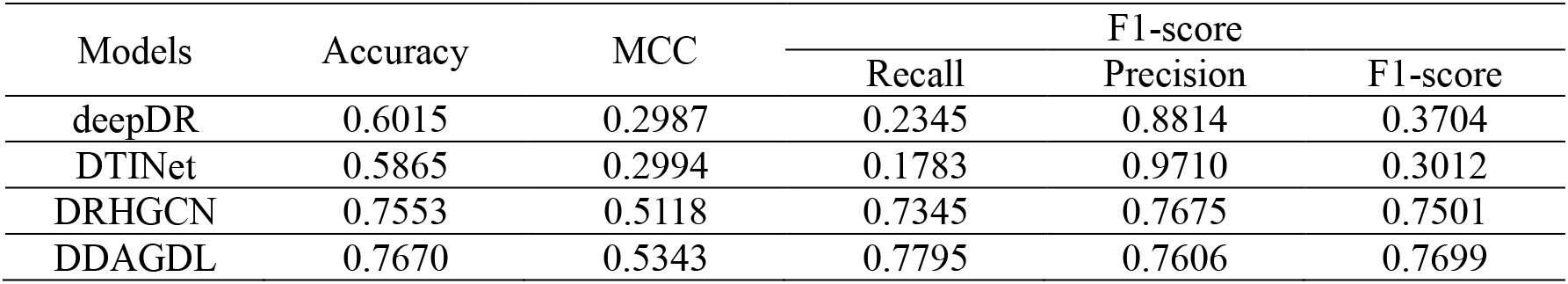
Experimental results of the various models under 10-fold CV on the B-dataset.

| Models | Accuracy | MCC | F1-score |  |  |
| --- | --- | --- | --- | --- | --- |
|  |  |  | Recall | Precision | F1-score |
| deepDR | 0.6015 | 0.2987 | 0.2345 | 0.8814 | 0.3704 |
| DTINet | 0.5865 | 0.2994 | 0.1783 | 0.9710 | 0.3012 |
| DRHGCN | 0.7553 | 0.5118 | 0.7345 | 0.7675 | 0.7501 |
| DDAGDL | 0.7670 | 0.5343 | 0.7795 | 0.7606 | 0.7699 |

**Table 2.** Experimental results of the various models under 10-fold CV on the C-dataset.

| Models | Accuracy | MCC | F1-score |  |  |
| --- | --- | --- | --- | --- | --- |
|  |  |  | Recall | Precision | F1-score |
| deepDR | 0.7696 | 0.6035 | 0.5450 | 0.9894 | 0.7022 |
| DTINet | 0.5683 | 0.2692 | 0.1370 | 0.9974 | 0.2401 |
| DRHGCN | 0.8124 | 0.6583 | 0.6552 | 0.9558 | 0.7772 |
| DDAGDL | 0.8420 | 0.6843 | 0.8499 | 0.8369 | 0.8432 |

**Table 3.** Experimental results of the various models under 10-fold CV on the F-dataset.

| Models | Accuracy | MCC | F1-score |  |  |
| --- | --- | --- | --- | --- | --- |
|  |  |  | Recall | Precision | F1-score |
| deepDR | 0.7501 | 0.5609 | 0.5241 | 0.9564 | 0.6762 |
| DTINet | 0.5420 | 0.2081 | 0.1118 | 0.1000 | 0.1545 |
| DRHGCN | 0.7783 | 0.5993 | 0.5938 | 0.9418 | 0.7279 |
| DDAGDL | 0.8443 | 0.6889 | 0.8520 | 0.8394 | 0.8455 |

**Figure 2.**
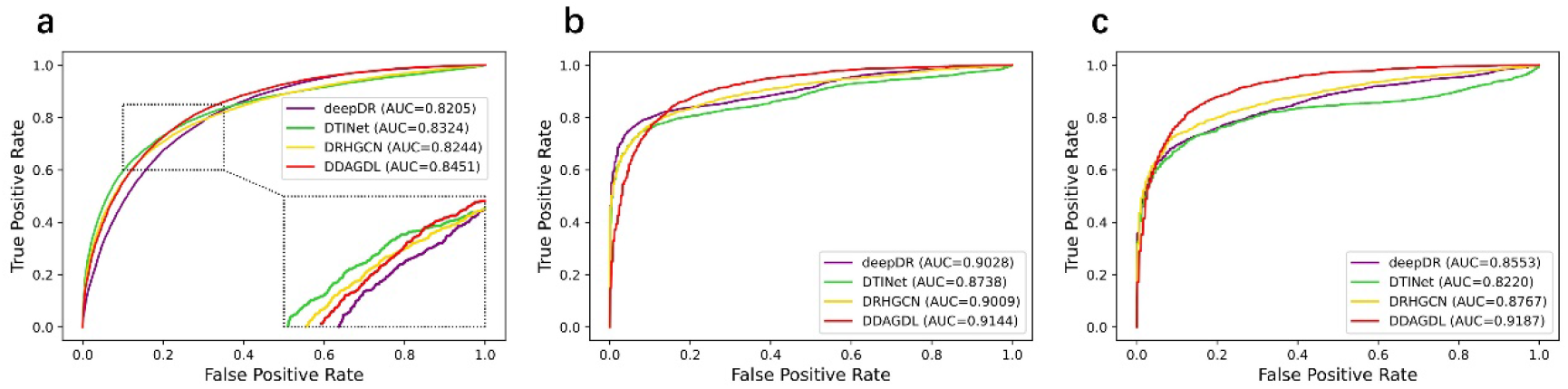
The ROC curves w.r.t. the overall performance of all comparing models on three benchmark datasets, and they are presented in subfigures (a)-(c), respectively.

In addition to its superior AUC, DDAGDL is also more robust than the other drug repositioning models as indicated by their evaluation scores. Take DTINet as an example, DDAGDL performs better by 25.22%, 37.69% and 58.76% than DTINet in terms of the average Accuracy, MCC and F1-score. In doing so, the distinguished performance of DDAGDL has demonstrated again, that it can adapt different datasets and achieve the best condition to infer new candidate compounds. We also note that DTINet has lower recall scores and higher precision scores, the main reason for that phenomenon is that over-fitting of the model results in an inability to accurately identify the true sample class. Similarly, deepDR and DRHGCN are for this reason. However, we need the model can provide more true positive samples as a useful reference for the drug research task, DDAGDL may have better advantages from a real demand perspective.

Although DDAGDL yields the best performance in terms of several evaluation metrics, an in-depth analysis is conducted of the results in Tables 1, 2 and 3 from another perspective. Regarding the lower performance of deepDR and DTINet for DDA prediction, the reasons accountable are as follows: (1) they need multiple types of drug-related network data to capture the features of drugs and diseases, and it is difficult to meet for a general dataset. (2) inferring unknown associations by relying on the similarity of the relationships between drugs and diseases, which ignores the role of molecular location information in the association network. (3) missing the biological signature of the molecule. In doing so, this approach makes it difficult to learn hidden information in HINs. DDAGDL not only takes into account the biological attributes, but also the geometric prior knowledge of network structure in a non-Euclidean domain is learned by the geometric deep learning strategy, its prediction ability is improved to better discover new DDAs in a more comprehensive manner. Moreover, DRHGCN achieves the second-best performance on all three benchmark datasets due to the fact that it uses GDL to mine drug and disease features, but suffers from over-smoothing and the resulting molecular features cannot be better expressed. In other words, the features learned will tend to be consistent, to lead the classifier to be difficult to distinguish. Hence, DDAGDL can better solve this flaw by multiple neural network propagation for each biomedical node when capturing their feature representations of them in a projected latent feature space.

In summary, these results indicated that considering the geometric prior knowledge of network structure in a non-Euclidean domain into DDA prediction is not a trivial task, while the geometric deep learning procedure of DDAGDL can simultaneously and effectively capture the underlying feature representations in the HIN, and further to improve the accuracy of DDA prediction.

### Non-Euclidean nature influence on the performance of DDAGDL

To better study the influence of the GDL strategy for drug repositioning, we have also constructed two variants of DDAGDL, i.e., DDAGDL-N and DDAGDL-A. In particular, DDAGDL-N merely contains network structure information, and its biological attributes are replaced by random Gaussian distribution initialization. DDAGDL-A is a model without network structure information, and simply uses biological attribute characteristics to train the prediction model. The XGBoost classifier with the same parameters as DDAGDL is applied to generate these two variant models, and then evaluated under 10-fold CV. The experimental results obtained from three benchmark datasets are presented in Tables 4, 5 and 6 and Figure 3, where several things can be noted. On the one hand, any variant cannot achieve desired performance in drug repositioning. In particular, DDAGDL performs better by 7.60% and 12.01% than DDAGDL-A and DDAGDL-N in terms of the average AUC across three benchmark datasets. One should that the evaluation metrics of DDAGDL-N are the lowest among DDAGDL’s variants. In this regard, only relying on the association network information may not be sufficient enough to accomplish the task of drug repositioning. On the other hand, DDAGDL-A shows a smaller margin in performance against DDAGDL-N in each evaluation metric. In particular, HINGRL-B performs better by 2.76%, 5.54%, 3.18%, 2.55% and 2.85% than HINGRL-A in terms of Accuracy, MCC, Recall, Precision and F1-score, respectively. This phenomenon suggests that the attributes of biomolecules are equally important as network structures, and should be taken into account when predicting the relationships between unknown drugs and diseases.

**Table 4.** Experimental results of performance comparison on the B-dataset.

| Models | Accuracy | MCC | F1-score |  |  |
| --- | --- | --- | --- | --- | --- |
|  |  |  | Recall | Precision | F1-score |
| DDAGDL-N | 0.7215 | 0.4430 | 0.7179 | 0.7231 | 0.7205 |
| DDAGDL-A | 0.7502 | 0.5004 | 0.7513 | 0.7497 | 0.7504 |
| DDAGDL | 0.7670 | 0.5343 | 0.7795 | 0.7606 | 0.7699 |

**Table 5.** Experimental results of performance comparison on the C-dataset.

| Models | Accuracy | MCC | F1-score |  |  |
| --- | --- | --- | --- | --- | --- |
|  |  |  | Recall | Precision | F1-score |
| DDAGDL-N | 0.7550 | 0.5101 | 0.7540 | 0.7559 | 0.7548 |
| DDAGDL-A | 0.7761 | 0.5526 | 0.7856 | 0.7713 | 0.7782 |
| DDAGDL | 0.8420 | 0.6843 | 0.8499 | 0.8369 | 0.8432 |

**Table 6.** Experimental results of performance comparison on the F-dataset.

| Models | Accuracy | MCC | F1-score |  |  |
| --- | --- | --- | --- | --- | --- |
|  |  |  | Recall | Precision | F1-score |
| DDAGDL-N | 0.6746 | 0.3495 | 0.6705 | 0.6760 | 0.6730 |
| DDAGDL-A | 0.7077 | 0.4160 | 0.7010 | 0.7106 | 0.7052 |
| DDAGDL | 0.8443 | 0.6889 | 0.8520 | 0.8394 | 0.8455 |

**Figure 3.**
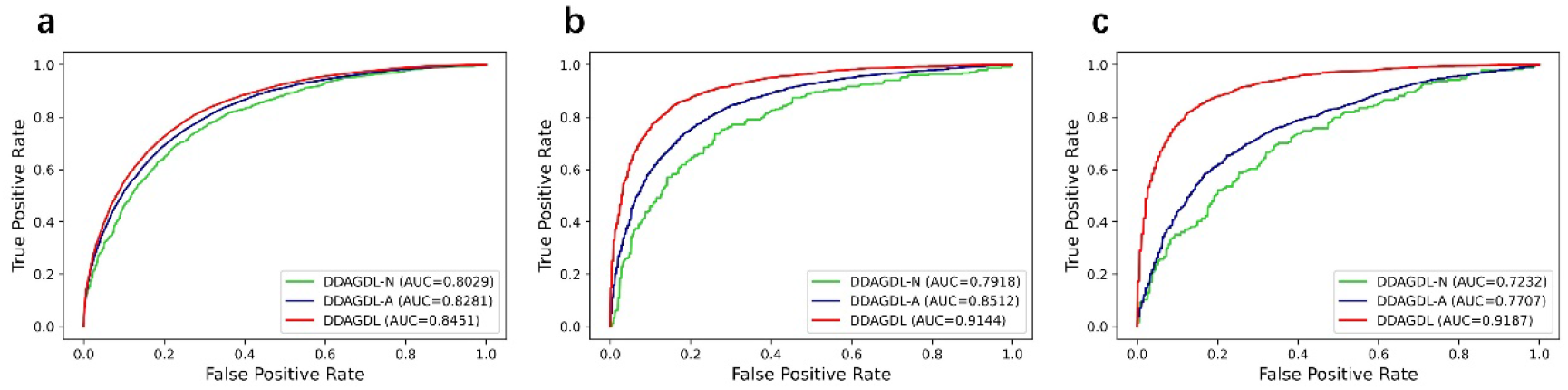
The ROC and PR curves are obtained by two variants of DDAGRL over three benchmark datasets in the ablation study, and they are presented in subfigures (a)-(d), respectively.

**Table 7.**
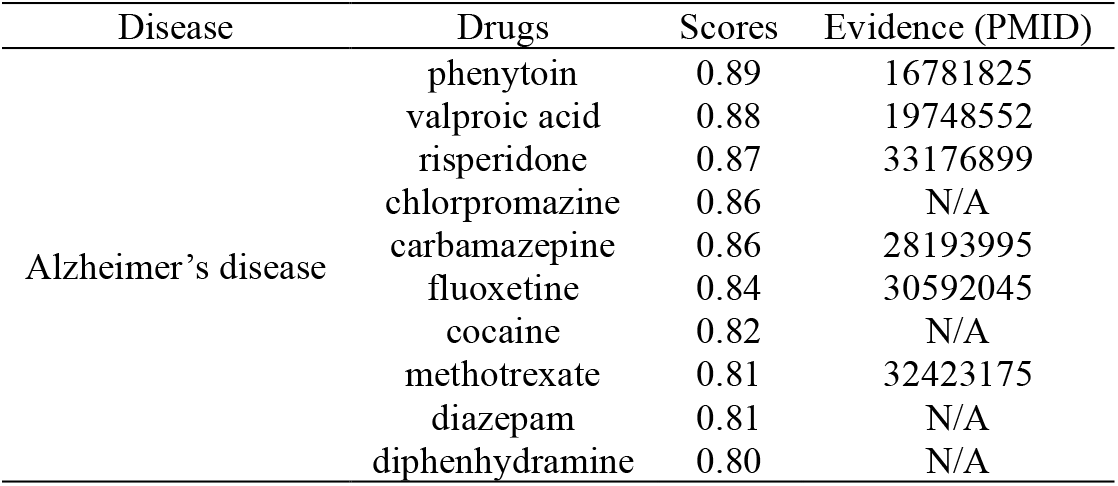
The top 10 candidate drugs predicted by DDAGDL for AD.

**Table 8.** The top 10 candidate drugs predicted by DDAGDL for BN.

| Disease | Drugs | Scores | Evidence (PMID) |
| --- | --- | --- | --- |
| Breast neoplasms | methylprednisolone | 0.94 | 12884026 |
|  | valproic acid | 0.92 | 30075223 |
|  | cocaine | 0.91 | N/A |
|  | nifedipine | 0.88 | N/A |
|  | phenytoin | 0.86 | 22678159 |
|  | simvastatin | 0.86 | 33705623 |
|  | amiodarone | 0.86 | N/A |
|  | sirolimus | 0.84 | 32335491 |
|  | ethinyl estradiol | 0.83 | N/A |
|  | betamethasone | 0.83 | N/A |

### Case studies

To demonstrate the capability of DDAGDL in practically discovering potential DDAs, we have conducted additional experiments on the B-dataset. In particular, all proven relationships between drugs and diseases are used to construct the training dataset and then DDAGDL is used to predict new candidate drugs for diseases. To delve into the experimental results of DDAGDL, we have presented the following two case studies for Alzheimer’s disease (AD) and Breast neoplasms (BN).

In table 4, the top 10 candidates discovered by DDAGDL for the potential treatment of AD, where 6 candidate compounds are evidenced in the relevant literature. Moreover, the candidate drugs for BN are also predicted by DDAGDL, and the top 10 results are presented in Table 5, of which 5 drug candidates are evidenced to be related to BN in the relevant literature.

To evaluate the superior performance of DDAGDL, we also have conducted the case studies on the other state-of-the-art models, the experiment results of which are shown in Supplementary material. In particular, the deepDR and DRHGCN models are selected to predict the candidate drugs for AD and BN on B-dataset. These experiment results of which are shown in Supplementary material. We note that deepDR and DRHGCN models have poor performance for discovering new candidate compounds. DRHGCN is the second-best model in the comparative experiment, only three candidates for AD are proved by the relevant literature in the top 10 results predicted, and eight candidates for BN fail to prove by the relevant literature. One should that the predicted scores of DRHGCN are lower than that of DDAGDL, which makes it difficult to accurately provide a reference for medical research. As a network-based model, when predicting AD drug candidates, only 2 of the top 10 results predicted by deepDR have been verified by relevant literature, while for BN’s drug candidate prediction, only one of them has been proved. One should that the predicted scores of deepDR are close to zero, which makes it difficult to discover new indications for approved drugs. Furthermore, we have performed an in-depth analysis of the experimental results from the perspective of a model designed. Regarding deepDR producing weak generalization ability, the reason accountable is that the model merely learns the features by relying on previous associations. For the disease to be predicted, the trained model is limited by whether the association of the input test is similar to the known association. Unlike deepDR, our model can convert the association data into a distinguishable space to obtain the features for drugs and diseases. The above analysis confirms that by using the GDL strategy to learn latent feature representations, DDAGDL can be a useful tool for drug repositioning due to its promising performance.

## Discussion

Drug repositioning is a promising strategy to discover new indicators of approved drugs, and thereby can improve traditional drug discovery and development, especially for previously untreated diseases. Recent advances in biomedical sciences, together with the development of artificial intelligence techniques, have further improved the effectiveness of computational drug repositioning approaches, which considerably facilitate the identification of top-ranked drug candidates by evaluating novel associations between drugs and diseases. In this work, we propose a new framework, namely DDAGDL, to predict DDAs by using geometric deep learning over a heterogeneous information network. More specifically, DDAGDL first integrates three kinds of drug-related networks, including drug-disease network, drug-protein network and protein-disease network, to compose a heterogeneous biomedical network, and a HIN is generated by further incorporating the biological knowledge of drugs, diseases and proteins. Second, DDAGDL makes use of complicated biological information to learn the feature representations of drugs and diseases with a geometric deep learning strategy, which allows DDAGDL to properly project drugs and diseases onto a latent feature space by additionally considering the geometric prior knowledge of network structure in a non-Euclidean domain. Finally, an XGBoost classifier is adopted by DDAGDL to complete the task of predicting DDAs. Experimental results demonstrate that DDAGRL yields a superior performance across all the three benchmark datasets under ten-fold cross-validation when compared with state-of-the-art prediction models in terms of several independent evaluation metrics. This could be a strong indicator that DDAGRL can effectively learn the feature representations of drugs and diseases by projecting complicated biological information, characterized by its non-Euclidean nature, onto a latent space with geometric deep learning. Hence, DDAGRL is capable of making full use of the geometric prior knowledge of HIN, and thereby enhancing the quality of feature representations of drugs and diseases. Furthermore, we have also conducted case studies to show the usefulness of DDAGRL in predicting novel DDAs by validating the top-ranked drug candidates predicted by DDAGRL for Alzheimer’s disease and Breast neoplasms. Our findings indicate that most of the drug candidates are of high quality, as they have already been reported by previously published studies, and some of them are not even found in the prediction results of the other comparing models. In this regard, leveraging geometric deep learning provides us an alternative view to address the problem of drug repurposing by properly handling the non-Euclidean nature of biomedical data, which has been ignored by most of the existing prediction models. In conclusion, we believe that our work opens a new avenue in drug repositioning with new insights gained from geometric deep learning.

In summary, the above experimental results have demonstrated the promising performance of DDAGDL in drug repositioning. On the one hand, DDAGDL simultaneously takes into account the attribute of biomolecules and network structures with non-Euclidean nature to obtain the feature representations of drugs and diseases. To be more specific, the traditional biological attributes have translation invariance in the Euclidean domain, which is limited by its lack of flexibility and weak expression ability, making it difficult to improve the accuracy of drug repositioning models. On this basis, we additionally consider the geometric prior knowledge of network structure in a non-Euclidean domain by making use of the GDL strategy to mine more underlying biologically meaningful characteristics, which further enhances the ability to express drug and disease features. On the other hand, DDAGDL improves the defects of existing GDL strategies. Specifically, DDAGDL first constructs the optimal projection space of each biomolecule by calculating the optimal number of propagation layers in neural networks, and then captures their feature representation from these spaces, which further improves the accuracy of our model in drug repositioning.

Although the experiment results have demonstrated the promising performance of DDAGDL, there are still some limitations to be addressed in the next work. First, known association network data comes from manually collected databases, which are easily introduced into noise to affect the training results. Therefore, we will construct a subgraph for each biomolecule to learn their representations. Second, we will introduce more types of association networks such as drug-drug association network [20] and drug-target association network [21], to enrich the HIN, from which DDAGDL is able to learn more expressive network representations of drugs and diseases.

## STAR Methods

### Datasets

To evaluate the performance of DDAGDL, three actual datasets are adopted to construct three HINs respectively, i.e., B-dataset, C-dataset and F-dataset. Each dataset contains three kinds of biological networks, i.e., drug-disease, drug-protein, and protein-disease networks. For B-dataset and F-dataset are collected from previous studies [16, 22, 23], in which B-dataset contains 18,416 DDAs, 3,110 drug-protein associations and 5,898 protein-disease associations, and F-dataset involves 1,933 DDAs, 3,243 drug-protein associations and 54,265 protein-disease associations. Moreover, C-dataset is also constructed by Luo et al.’s instruction [24], it contains 2,532 DDAs, 3,773 drug-protein associations and 10,734 protein-disease associations. The drug-protein associations and protein-disease associations are downloaded from the DrugBank database [25] and the DisGeNET database [26], respectively.

### Construction of HIN

To better describe the procedure of DDAGDL, we have introduced a three-element tuple, i.e. HIN(***V, C, EE***), where ***V*** = {*V*^*dr*^, *V*^*pr*^, *V*^*di*^} is a set of drugs (*V*^*dr*^), proteins (*V*^*pr*^), diseases (*V*^*di*^) that are involved to construct a HIN, ***E*** = {*E*^*dd*^, *E*^*dp*^, *E*^*pd*^} represents the drug-disease network (*E*^*dd*^), the drug-protein network (*E*^*dp*^), the protein-disease network (*E*^*pd*^), ***C*** = [*C*^*dr*^; *C*^*pr*^; *C*^*di*^]^*T*^ ∈ *ℝ*^|***V***|×*d*^ denotes the calculated biological attributes for all nodes in HIN, where |***V***| is the number of all nodes. Moreover, *N* and *M* are used to denote the number of drugs and diseases, the adjacency matrix of HIN is defined as *A* ∈ *ℝ*^|***V***|×|***V***|^.

### Calculating biological attributes

Regarding the biological attributes for drugs, diseases and proteins, three different computer algorithms are used. We collect three kinds of biological knowledge, i.e., the Simplified Molecular Input Line Entry System (SMILES) [27] for drugs, the sequence information of proteins, and Medical Subject heading (MeSH) descriptors of diseases. In addition, disease biological attributes based on disease phenotype by using MimMiner [28], and drug biological attributes based on chemical structures [24] are used as an alternative when using C-dataset, respectively.

To facilitate calculation when geometric deep learning, we first have performed the RDkit toolkit [29] to obtain the biological attributes *C*^*dr*^ by calculating the SMILES of drugs. Second, the biological attributes *C*^*di*^ are obtained based on the MeSH descriptors by Guo et al.’s instruction [30]. After that, the sequence information of proteins is divided into four classes according to the nature of the side chain, i.e., (Ala, Val, Leu, Ile, Met, Phe, Trp, Pro), (Gly, Ser, Thr, Cys, Asn, Gln, Tyr), (Arg, Lys, His) and (Asp, Glu), and then a 3-mer algorithm [31-35] is used to obtain the biological attributes *C*^*pr*^. Finally, all biological attributes ***C*** obtained are unified into 64-dimension by an autoencoder scheme [36].

### Extracting feature representations

Traditional deep learning cannot effectively learn non-Euclidean data, such as biological gene protein data, chemical composition structure data and biological association network data. The recent rise of geometric deep learning makes it easier to study the associations between biological entities. In order to better meet our research problem, we design a geometric deep learning algorithm based on a graph convolutional neural network to extract feature representation for drugs and diseases, which can project the geometric prior knowledge of network structure with non-Euclidean data into a latent feature space to obtain a more influential of feature representations for drugs and diseases. In particular, a general graph convolutional neural network [37] is defined as:

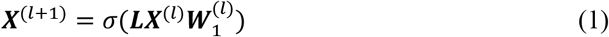

Where 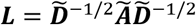 denotes the normalized Laplacian matrix, ***Ã***; = ***A*** + *I* is an adjacency matrix with added self-loops 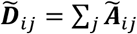 represents a degree matrix. ***W***_*1*_ as a random weight matrix, *σ*(.) is an activation function and *l* represents the number of layers for the neural network. In this work, we define the first layer ***X***^(0)^ = ***C***, and assume ***W***_*1*_ is the identity matrix and *σ*(.) is an identity function according to Wu et al. [38] and Li et al. [39]. Hence, Eq. (1) can be reconstructed as the following form.

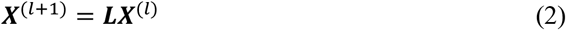

When the neural network layer *l* is large enough [40]. the Eq. (2) is considered as:

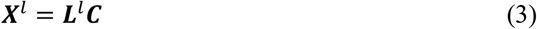

where *L*^*l*^ is the change matrix after more than one layer in neural networks. In other words, the operation of multiple neural network propagations can be regarded as the transformation form of the biological feature matrix ***C***.

In order to make up for the inherent over-smoothness caused by the defect of the graph convolutional network, we calculate the optimal number of neural network propagation layers for each biomedical entity to better learn their representation. Let us introduce a function *K* to calculate the optimal number of the layer *l* for node *v*_*i*_ (*v*_*i*_ ∈ ***V***).

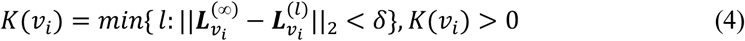

where *δ* is a parameter (*δ* > 0), and ||. ||_2_ denotes the function with two-norm. To better deal with the features after multiple neural network propagation, we design an attention function to aggregate *l* kinds of features. After that, Eq. (3) can be translated as follow.

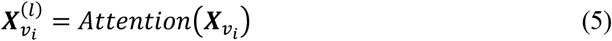

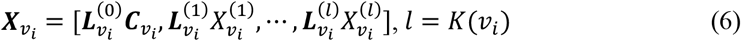

Moreover, we add the initial feature, i.e. the biological feature ***C***, to enhance the expression of features in the course of the multiple neural network propagation. For instance,

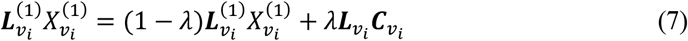

Regarding the attention function, its function is as a pooling layer to aggregate all features, the details as:

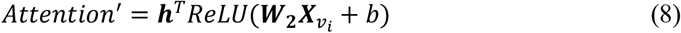

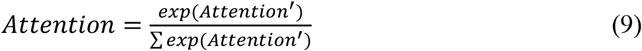

where ***W***_2_ is a *l* × *l* weight matrix, *b* is bias and ***h*** is a parameter according to Xiao et al.’ description [41]. At last, a (*N* + *M*) × *d* matrix ***X*** is constructed to denote the feature representations of drugs and diseases.

### Identifying new DDAs

After extracting the feature representations of drugs and diseases from the projected feature space, DDAGDL next aims to predict the relationships between drugs and diseases on the base of their learned representations. In particular, we first use a typical machine learning classifier, i.e. XGBoost [42], to complete the task of DDA prediction. Then, we compose a set of drug-disease pairs denoted as *F* = {(*F*_*i*_, *y*_*i*_)}(1 ≤ *i* ≤ |*F*|), where *F*_*i*_ denote the concatenated feature vector of the *i*-th drug-disease pair, *y*_*i*_ ∈ {0,1} represent the label of this pair and the value of *y*_*i*_ is 1 if connected and 0 otherwise. *F*_*i*_ is the concatenation of ***X***^***dr***^ and ***X***^***d****i*^, which are the respective representation vectors of drug 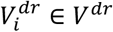 and disease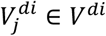. Furthermore, a result matrix ***R*** is introduce to collect the prediction scores between drugs and diseases whose associations are unknown in advance.

## Notes

### Competing Interest Statement

The authors have declared no competing interest.

